# An Anterior Cingulate Cortex Neuronal Ensemble Controls Contextual Opioid Analgesic Tolerance

**DOI:** 10.1101/2025.08.16.670663

**Authors:** Rafael E. Perez, Yann S. Mineur, Cheryl Chen, Wen-Liang Zhou, Julian L. Thompson, Saagar S. Motupally, Angélica Minier-Toribio, Marina R. Picciotto

## Abstract

Opioids are first-line treatments for pain; however, tolerance leads to decreased efficacy and escalated dosing, increasing the risk of opioid use disorder and overdose. Despite well-established cellular and molecular adaptations, opioid analgesic tolerance can be rapidly reversed in settings where these drugs are not expected. The specific neuronal populations that orchestrate this expectation-based tolerance are not known. In this study, we used a contextual tolerance training method, whole-brain clearing, and immunostaining to identify brain regions involved in contextual tolerance and to pinpoint a neuronal ensemble in the anterior cingulate cortex (ACC) that is activated by contextual analgesic tolerance. We observed that calcium activity in principal neurons of the ACC is suppressed by fentanyl in opioid-naïve mice and during contextual reversal, but not during contextual tolerance. Chemogenetic silencing of the ACC induced analgesic tolerance reversal in the opioid-associated context without affecting thermal nociception in opioid-free mice. Conversely, chemogenetic activation of the ACC contextual tolerance-active neuronal ensemble triggered analgesic tolerance in an unassociated context. This research highlights the role of ACC neuronal ensembles in mediating expectation-driven, contextual opioid analgesic tolerance without affecting basal nociception. Modulating ACC activity could provide a promising strategy to improve pain relief while maintaining the essential ability to detect harmful stimuli.

## Introduction

Opioid drugs remain critical for clinical pain management (1), however opioid overdoses are one of the leading causes of accidental deaths in the United States(2). Chronic use of these medications leads to tolerance to their analgesic effects, requiring higher doses to manage pain effectively(3, 4). As doses used to alleviate pain increase, they can reach dangerous levels that may result in fatal respiratory depression. Thus, this escalation in dosage significantly increases the risk of opioid misuse and opioid use disorders, contributing to the dramatic rise in overdose deaths reported over the last 30 years (5).

Extensive research has focused on the molecular and pharmacological mechanisms underlying opioid analgesic tolerance, including receptor desensitization and trafficking(6). However, environmental contexts also influence the development of analgesic tolerance significantly. When opioids are used repeatedly in specific contexts (e.g., home vs. hospital), tolerance develops selectively within those environments(7). Critically, opioid use in unfamiliar settings can lead to loss of this contextual tolerance, contributing to overdoses when patients take their usual dose in novel environments, a phenomenon documented in overdose case reports(7–9). Although contextual tolerance reversal can have important consequences for analgesia and overdose, the biological mechanisms underlying this phenomenon remain poorly defined. Further investigation into the brain regions involved in the environmental regulation of opioid analgesic tolerance is essential to develop better therapeutic strategies for the treatment of pain that minimize use and addiction risk.

Contextual opioid analgesic tolerance is a learned response in which contextual cues are associated with, and come to predict, the effects of the drug, thereby modifying nociception. This suggests that the mechanisms underlying contextual tolerance likely involve brain regions that integrate contextual learning, expectation, nociception, and analgesia. One such area is the anterior cingulate cortex (ACC), a central site of opioid analgesic action that plays a critical role in environmental modulation of pain and analgesia (10–14). Moreover, ACC dysfunction is implicated in chronic pain states and opioid use disorders, making it a clinically relevant therapeutic target(15–19). ACC activity predicts pain severity, and deep-brain stimulation in this region alleviates chronic pain in patients(20). Additionally, the ACC is vital for contextual memory formation, with ACC neuronal ensembles encoding distinct contextual features through direct projections to the hippocampus, thereby facilitating memory formation and recall(21, 22). Collectively, these findings indicate that the ACC may serve as a pivotal regulator of contextual tolerance.

Here, we identify an ACC neuronal ensemble as a critical mediator of contextual opioid analgesic tolerance to acute thermal stimuli using a robust conditioning paradigm, targeted recombination in active populations (TRAP) of neurons, neural Ca^2+^ imaging in behaving mice, and chemogenetic manipulations. Our findings also highlight the modulation of ACC activity as a strategy for rapidly reversing analgesic tolerance in drug-associated environments. This positions ACC-targeted interventions as a potential therapeutic approach to enhance pain relief, particularly in opioid-dependent populations, while reducing tolerance and overdose risk.

## Results

To investigate the mechanisms driving contextual tolerance, we established a behavioral model that mimics expectation-based contextual opioid analgesic tolerance. Mice received daily injections of fentanyl (25 µg/kg) in one context, alternating every other day with saline injections in a different context (**Fig. 1A**). Fentanyl initially increased latency to display nociceptive behaviors in the hot plate test. This analgesic response decreased by 50% after 7 fentanyl sessions, indicating the development of tolerance (**Fig. 1B**). We then tested whether tolerance was dependent on the expectation of the drug in the context by injecting mice with fentanyl and placing them in the saline-associated context or a new context. We found that placement in the saline-associated context or the new context before hotplate exposure significantly reversed tolerance in male mice, with a strong trend towards reversal in female mice (**Fig. 1C, sFig 2**). Thus, this paradigm reliably induces reversible context-dependent tolerance in male animals and is sensitive to the expectation of outcome before exposure to a nociceptive experience.

**Figure 1.**
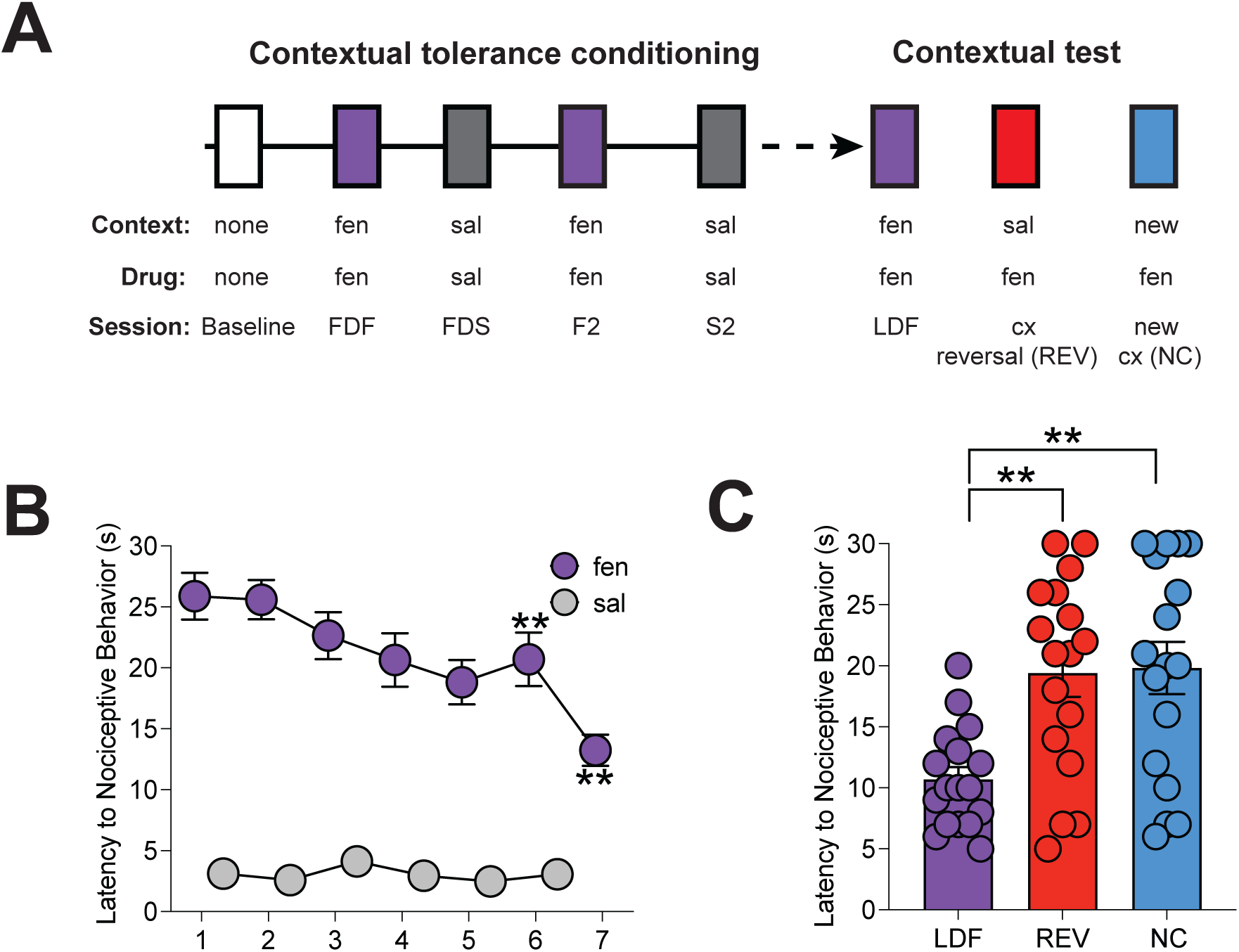
A conditioning procedure that induces contextual opioid analgesic tolerance in male mice. **A.** Experimental design: Mice received alternating daily injections of fentanyl (fen) or saline (sal) for 14 days in distinct contexts (cx), followed by daily assessment of antinociceptive response on a hotplate at 56°C. After conditioning, contextual tolerance was tested by administering fen in the previously sal-paired or a new cx and evaluating nociceptive latencies. **B.** A decrease in the antinociceptive effect of fen was observed over time, with reductions notable during sessions 5 and 7 when compared to the first fen session (FDF). No changes in nociception were observed after repeated sal injections. **C.** Mice showed a statistically significant reversal of analgesic tolerance compared to the Last Day of Fentanyl administration (LDF) in the sal-paired context (REV) or in a new context (NC) (n = 17).

Next, to identify cells active after re-exposure to either the fentanyl-(tolerance) or saline-paired (reversal) context, we marked brain-wide ensembles of neurons activated during contextual tolerance and its reversal in ArcCreER:Ai14 (ArcTRAP:tdTomato) mice (23), and used cFos labeling following tissue clearing. We used a modified engram index, previously applied to whole-brain cFos data, to identify candidate regions with increased activity following re-exposure to the fentanyl context for further analysis (**Fig. 2A**) (24). Exposure to the fentanyl-associated context increased the fentanyl-context engram index of brain areas known to be involved in memory, opioid hyperalgesia, and physiological homeostasis (**Fig. 2B**, **Supplementary Table 1**)(16, 25–27). Interestingly, <u>tolerance reversal</u> increases neuronal activity in brain regions implicated in environmental novelty detection and contextual memory suppression, such as the perirhinal cortex (**sFig. 3C, Supplementary Table 2**)(28, 29). These findings support the accuracy of our whole-brain activity tagging results. Beyond identifying brain regions involved in contextual tolerance, these findings also suggest that reversing tolerance by changing context likely relies on an active process that recruits brain regions involved in novelty detection (30).

**Figure 2.**
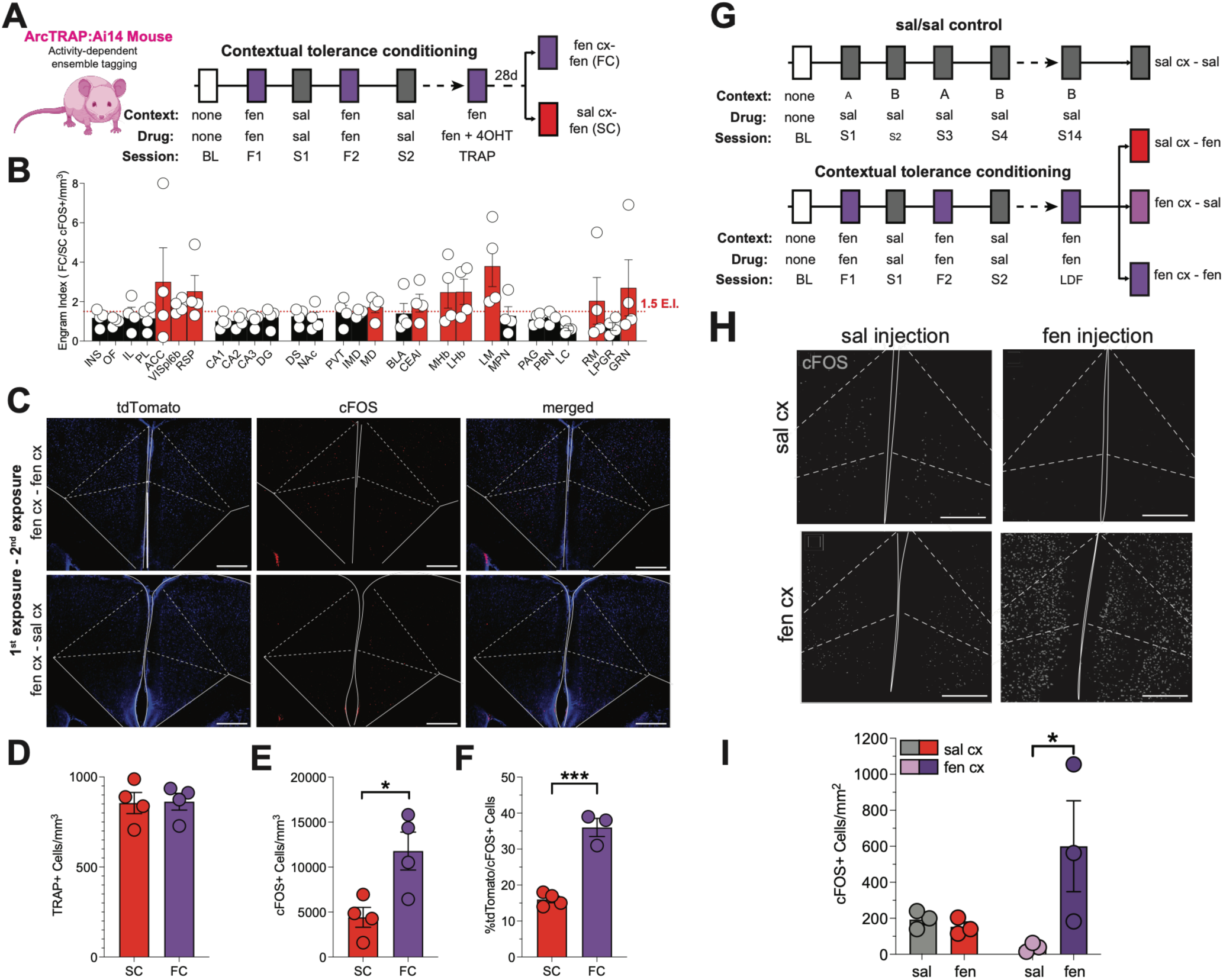
Contextual opioid tolerance induces brain-wide activity changes and the recruitment of an ACC neuronal ensemble. **A.** Experimental design for whole-brain activity mapping: ArcTRAP: Ai14 mice were administered 4-hydroxytamoxifen (4OHT, 50 mg/kg) on the last day of fen administration in the fen-paired cx (TRAP). Four weeks later, mice were re-exposed to fen in either the fen (FC) or sal cx (SC). **B.** Brain regions with increased ratio of cFos/tdTomato expression (FC / TRAP) following re-exposure to fen in the previously fen-paired cx. **C.** Representative images showing tdTomato+ (TRAP) and cFos+ cells in the ACC (scale bar = 400μm). **D.** There was no difference in the number of TRAP cells across groups, consistent with equivalent fen and cx exposure in all mice during initial ensemble trapping. **E-F.** There was an increase in cFos+ and cFos+/TRAP cells in the ACC of mice re-exposed to fen in the previously fen-paired cx (n = 4 per group). **G**. Experimental design for evaluation of cFos labeling in response to cx, fen, or the combination of cx + fen. Control mice received sal in alternating cxs for 14 days (S1-S14), and experimental animals underwent fentanyl contextual tolerance conditioning before exposure to the previously fen-paired cx alone, fen alone, or the fen-paired cx + fen. **H.** Representative microscopy images showing cFos+ cells in ACC (scale bar = 400μm). **I.** There was a significant increase in cFos+ cells in ACC following fen administration in the fen-paired cx (n = 3 per group; *p < .05, **p < .01, ****p < .0001). Data are shown as means ± SEM and individual data points. INS = insular cortex, OF = orbitofrontal cortex, IL = infralimbic cortex, PL = prelimbic cortex, VISpl6b = posterolateral visual cortex layer 6b, RSP = retrosplenial cortex, CA1 = hippocampal area CA1, CA2 = hippocampal are CA2, DG = hippocampal area dentate gyrus, DS = dorsal striatum, NAc = nucleus accumbens, PVT = paraventricular nucleus of the thalamus, IMD = interomediodorsal thalamus, MD = mediodorsal thalamus, BLA = basolateral amygdala, CEAl = lateral central nucleus of the amygdala, MHb = medial habenula, LHb = lateral habenula, LM = Lateral mammillary nucleus, MPN = medial preoptic nucleus, PAG = periaqueductal gray, PBN = parabrachial nucleus, LC = locus coeruleus, RM = nucleus raphe magnus, LPGR = lateral paragigantocellular nucleus, GRN = Gigantocellular reticular nucleus.

We confirmed the whole-brain study findings using cFos labeling in candidate regions with 4 conditions controlling for independent effects of drug and context alone (**Fig. 2G, sFig. 3**). In both studies, the ACC was highly activated only when fentanyl was administered in the fentanyl-paired context (**Fig. 2E, I**). In addition, there was 40% colocalization between ACC cells tagged on the last day of fentanyl training (tdTomato) and those labeled after re-exposure to fentanyl in the fentanyl context (cFos) four weeks later, suggesting that these cells could represent a persistent tolerance-sensing neuronal ensemble (**Fig. 2D-F**).

To evaluate the role of the ACC in the development and expression of contextual tolerance, we used *in vivo* fiber photometry to measure calcium transients with the calcium reporter GCamP6 (AAV5-CaMK2-GCaMP6f) as an indicator of neuronal activity in ACC neurons of behaving mice (31). Fluorescent signals were measured during contextual exposure throughout the conditioning procedure and reversal testing (**Fig. 3C-H**). Compared to the baseline session or subsequent saline sessions (**Fig. 3C-E**), the first fentanyl administration initially suppressed ACC activity, then caused a rebound in activity exceeding baseline levels several minutes later during context exposure (**Fig. 3F**), consistent with previous findings (32). By the 7th fentanyl training session, the early suppression of ACC activity and the late increase in activity were lessened, indicating the development of tolerance to the modulatory effects of fentanyl (**Fig. 3G**). Both the early and late effects of fentanyl were restored when fentanyl was administered in the previously saline-paired context (**Fig. 3H**). These photometric patterns correlate with the levels of fentanyl-induced analgesia and tolerance (**Fig. 3K**), with a significant negative correlation observed between the magnitude of the early decrease in signal within the context and the nociception response in the hotplate (**Fig. 3L**). Conversely, a positive correlation was found between the late increase in signal within the context and nociception (**Fig. 3M**).

**Figure 3.**
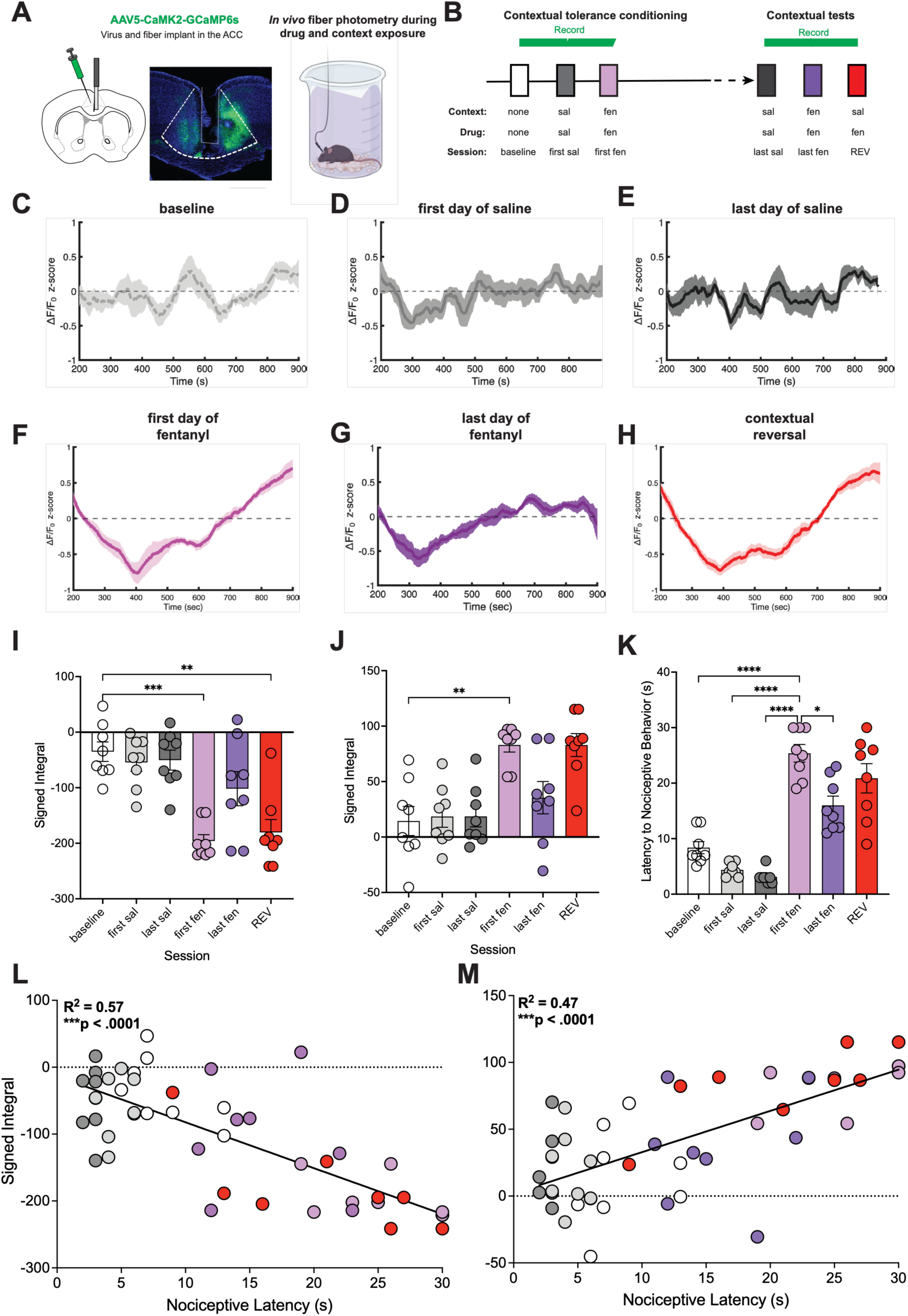
ACC activity in the fentanyl-paired context is correlated with fentanyl-induced antinociception across the contextual tolerance procedure and during tolerance reversal. **A.** Schematic and representative image illustrating expression of the calcium indicator GCaMP6s and the implanted fiber in the anterior cingulate cortex (ACC). **B.** Timeline of experimental procedures. Each day, mice were administered either sal or fen, connected to the fiber patch cord, and placed in the sal- or fen-paired cx for 15 minutes before nociceptive testing on the hotplate. **C-H.** Photometric trace depicting ACC calcium signals. **C.** Baseline recording day with no injection. **D.** First day of saline administration. **E.** Last day of saline administration. **F.** First day of fentanyl administration. **G.** Last day of fentanyl administration. **H.** Contextual reversal (REV, fen administered in the sal-paired cx). **I, J.** Quantification of early and late phases of photometric responses. **I.** There was a significant decrease in the signed integral of the early (200-650s) photometric signal during both the first day of fentanyl administration and REV when compared to baseline. **J.** Conversely, there was a significant increase in the signed integral of the photometric signal during the late phase (650-900s) of context exposure during the first day of fentanyl administration. **L.** A negative correlation was found between the calcium signal during the early phase of fentanyl-paired context exposure and nociceptive behaviors. **M.** A positive correlation was identified between the calcium signal during the late part of the contextual exposure and nociceptive behaviors, n = 8 (*p < .05, **p < 0.01, ***p < 0.001. Data displayed as means ± SEM and individual data points.

In contrast to changes in calcium activity during contextual exposure, we observed increases in ACC activity after hotplate placement during the baseline and subsequent saline sessions (**Fig. 4A-C**). This effect was completely blunted following initial fentanyl administration (**Fig. 4D**), with no tolerance development after 7 fentanyl/context pairings or a fentanyl/saline context pairing (**Fig. 4E**). We also found a significant inverse correlation between nociception-evoked ACC activity and hotplate latencies, consistent with previous reports (33). These findings indicate that ACC activity during hotplate exposure is unaffected by contextual tolerance or reversal and align with recent reports showing that nociception-evoked ACC activity does not vary with contextual training (34). Together, the findings from photometry experiments indicate that anticipatory ACC activity during **context exposure** is key for contextual analgesic tolerance and its reversal.

**Figure 4.**
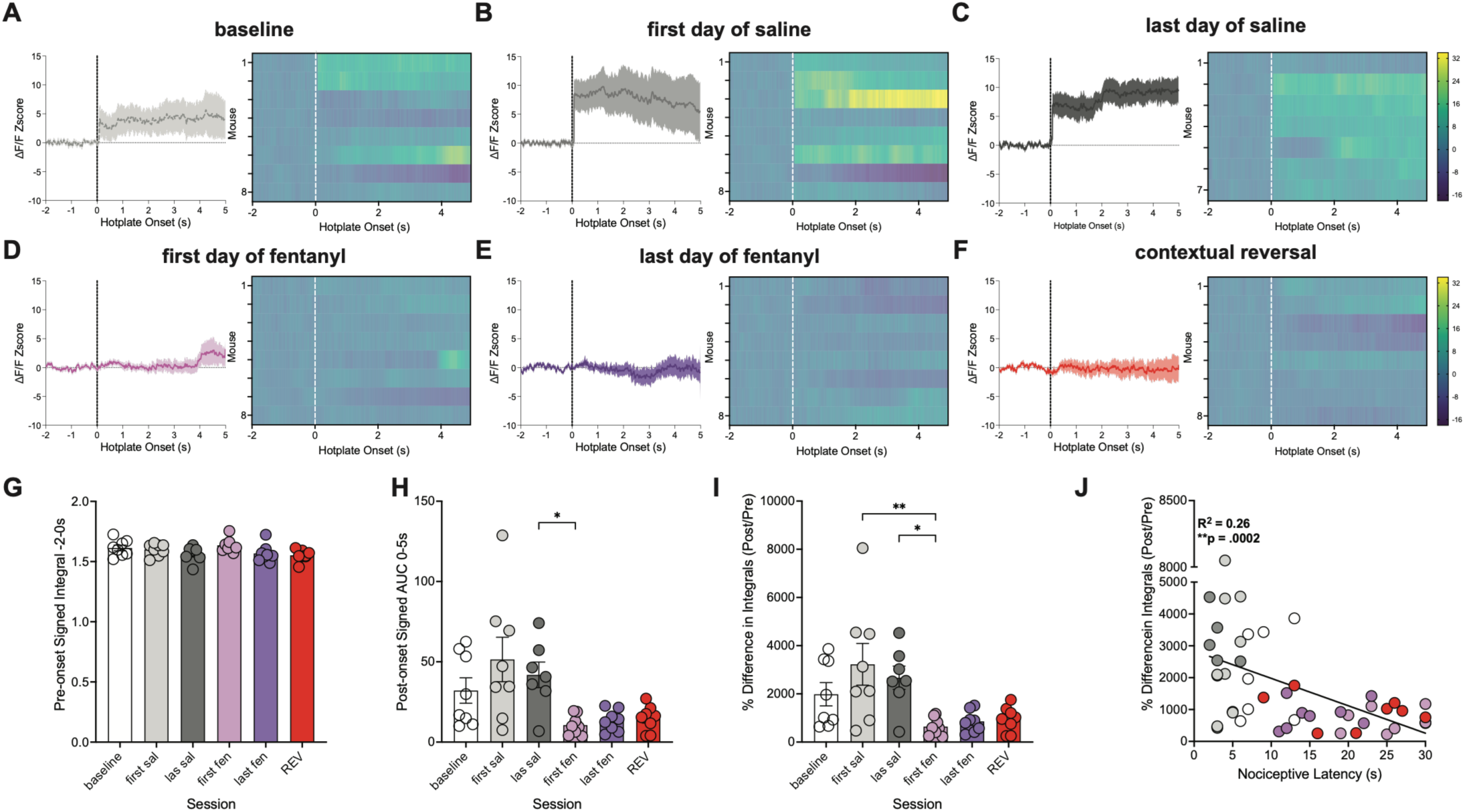
Hotplate-induced ACC activity increases are suppressed by fentanyl and are not susceptible to contextual tolerance. A-C. Hotplate placement increases ACC activity at baseline, on the first day of saline, and on the last day of saline fentanyl-paired context exposure. **D-F.** Fentanyl administration blocks hotplate-induced increases in ACC activity after the first and last exposures to the fentanyl-paired context, and after contextual reversal. **G** Signed integral values during the 2 sec prior to hotplate placement. **H.** Signed integral values during the 0-5s period post hotplate placement. There is a significant decrease in signal after the first fentanyl-paired context exposure when compared to the last saline-paired context exposure session. **I.** There is a significant decrease in the ratio of the signal (Post/Pre) after the first fentanyl-paired context exposure when compared to the first and last saline-paired context exposure sessions. **J.** A negative correlation was found between the ratio of the signal pre- and post hotplate placement and nociceptive behaviors, n = 8 (*p < .05, **p < 0.01. Data displayed as means ± SEM and individual data points.

Finally, to assess whether ACC activity and the context- and fentanyl-active ACC neuronal ensemble are necessary and sufficient for contextual analgesic tolerance, we employed a chemogenetic method to inhibit the ACC pan-neuronally (Gi-DREADD, AAV5-hsyn-hM4Di-mcherry) or selectively activate the contextual tolerance neuronal ensemble (AAV2-DIO-hM3Gq-mCitrine in ArcTRAP mice) (**Fig. 5A, B**). Administration of the DREADD ligand clozapine-N-oxide (CNO) did not alter nociception in response to saline administration or when fentanyl was administered to control mice. However, CNO reversed tolerance in Gi-DREADD-expressing mice when fentanyl was administered in the fentanyl-paired context (**Fig. 5C**). Thus, inhibiting ACC activity blocks contextual tolerance without altering nociception.

**Figure 5.**
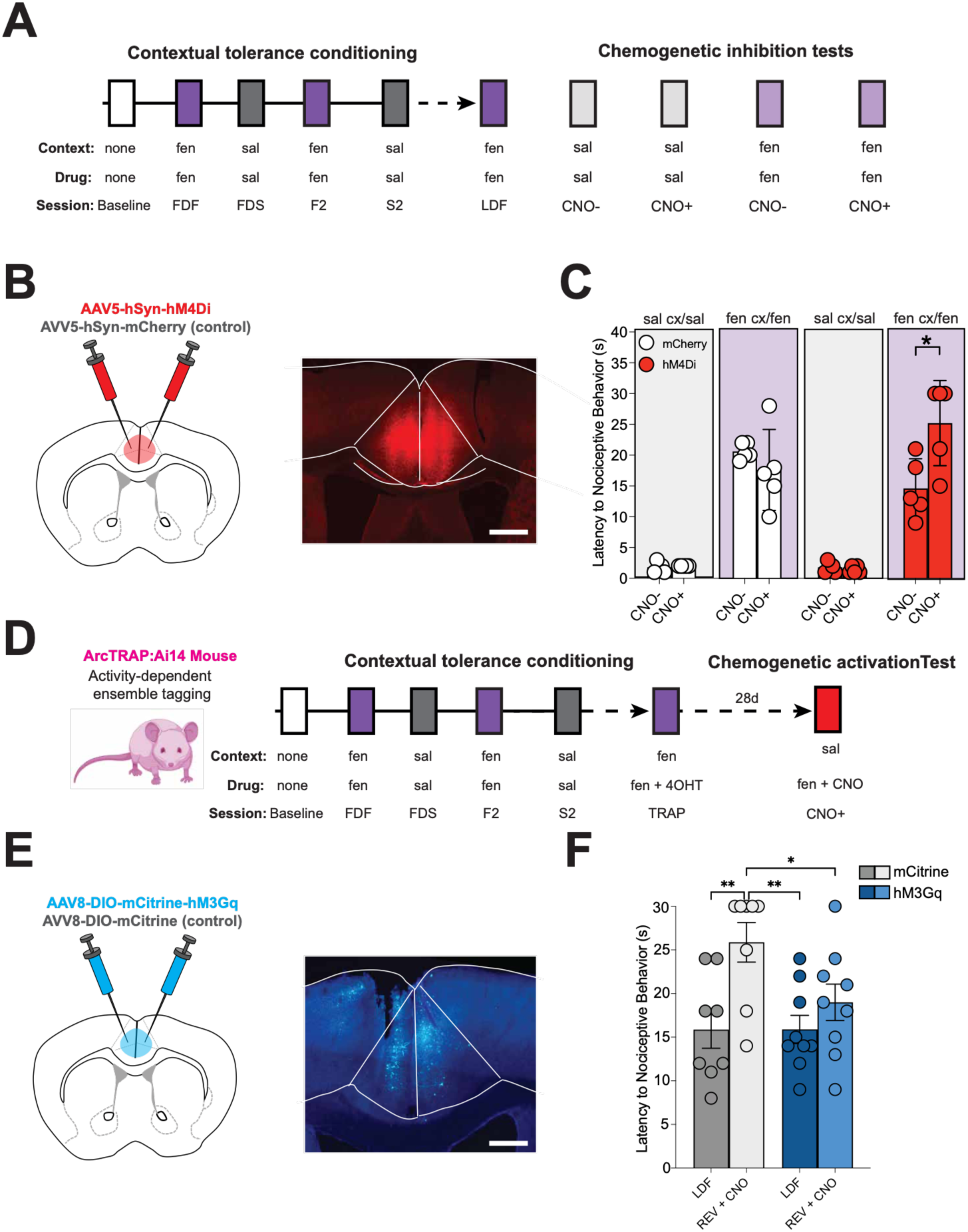
ACC tolerance-active ensemble neurons mediate contextual tolerance to fentanyl analgesia. **A.** Timeline and experimental design for the chemogenetic inhibition experiment. After mice underwent contextual analgesic tolerance training, they were exposed to either sal in the sal-paired cx or fen in the fen-paired cx, with or without the chemogenetic ligand clozapine-N-oxide (CNO). **B.** Schematic and representative image of viral injection of hsyn-hM4Di (hM4Di) in the ACC (scale bar = 500μm). **C.** Administration of CNO had no effect on nociceptive responses on the last day of sal administration in the sal cx in control or hM4Di-expressing mice, or fen administration in the fen cx in control mice, but resulted in increased nociceptive latencies in hM4Di-expressing mice following the last day of fen exposure in the fen cx, n = 5 per group. **D.** Experimental timeline. Mice underwent contextual analgesic tolerance conditioning until they developed tolerance, and on the last day of fen administration, were injected with the TRAP ligand 4OHT to induce CRE expression in active cells. Four weeks after 4-OHT administration, mice were placed in the sal-paired context and administered CNO and fentanyl (TRAP). **E.** Schematic of strategy to stimulate tolerance-active ensembles in the ACC. Male ArcTRAP mice were injected with viruses containing a cre-dependent chemogenetic activator, hM3Gq, or an mCitrine control. Representative image showing hM3Gq expression in the ACC (scale bar = 500μm). **G.** CNO administration had no effect in control mice but prevented contextual tolerance reversal in hM3Gq-expressing mice, n = 8 or 9 per group (*p < 0.05, **p < .01). Data displayed as means +/- SEM and individual data points.

We next tested whether reactivating ACC tolerance-active ensemble neurons is sufficient to induce tolerance in an unassociated context (**Fig. 5D, E**). Ensemble neurons were tagged on the last day of fentanyl exposure as described above, with one notable exception: mice were not placed on the hot plate after fentanyl and contextual exposure to avoid labeling ACC cells that encode nociception (see **Fig. 4**) (23, 35). Three weeks after ensemble labeling, mice were administered fentanyl and CNO in the previously saline-paired context (**Fig. 5F**). As expected, control mice showed reversal of analgesic tolerance after exposure to the saline-paired context. However, excitation of ACC-tagged ensembles in Gq-DREADD-expressing mice with CNO was sufficient to maintain fentanyl tolerance (prevent reversal) in the saline-paired environment (**Fig. 5F**). Therefore, selective modulation of ACC neuronal populations can either induce or prevent contextual analgesic tolerance without affecting nociception.

## Discussion

In this study, using a novel conditioning procedure in mice, we show that ACC activity encodes anticipation of drug-analgesic responses to thermal stimuli in an associated context, thereby inducing tolerance that can be quickly reversed despite profound molecular and cellular adaptations (6). We used this paradigm to show that the activity of ACC neurons tracks the development and expression of contextual opioid tolerance. Consistent with these changes in ACC, chemogenetic regulation of ACC neurons or the specific neuronal ensemble activated by contextual exposure is sufficient for contextual tolerance and necessary for its reversal.

ACC regulation of contextual analgesic tolerance is specific to the administration of opioids in the opioid-paired context, a condition in which the drug is expected. These results align with previous work showing the ACC plays a crucial role in regulating anticipated pain and analgesia(13, 36, 37). In contrast to a recent study that compared morphine analgesic tolerance in mice receiving repeated escalating doses of morphine either in their home cage or in a different context (Hou *et al*)(38), the current experiments identify the ACC as a critical mediator of tolerance in a paradigm that measures expectancy as an important parameter in both contextual tolerance and its reversal(38).

The current findings align with previous research showing that ACC is necessary for learned expectations of pain, pain avoidance, and pain relief(13, 37, 39, 40). These results build on earlier work by being the first to identify the role of the ACC in expectation-based analgesic tolerance, independent of pain modulation. We found that ACC activity increases only when <u>both</u> context and drug are presented together, and not after saline treatment, the condition with the highest degree of nociception. These *in vivo* photometric results also reveal a discrepancy in how contextual training influences ACC activity during exposure to the <u>context</u> compared to hotplate testing. Intriguingly, repeated contextual exposure suppressed the effects of fentanyl while animals were in the drug-paired context, but did not change fentanyl’s effects during hotplate testing. These findings point towards a dissociation between the ACC’s role in contextual and nociceptive processing.

Further, chemogenetic silencing of ACC activity did not change nociceptive responses in mice treated with saline. Notably, while some studies have reported effects of ACC manipulation on basal nociception in various pain paradigms, particularly chronic pain assays(18, 19), we did not observe such effects in the acute thermal hotplate test. This discrepancy likely reflects differences in pain modality, testing conditions, and contextual factors across experimental paradigms, highlighting the importance of considering environmental and methodological variables when interpreting ACC’s role in nociception. Future studies systematically comparing ACC function across different pain modalities and contextual conditions will be important to resolve these apparent inconsistencies. Additionally, stimulating an ACC fentanyl-context active neuronal ensemble tagged in the absence of a pain experience (no hotplate exposure) is sufficient to prevent reversal of acute thermal analgesic tolerance in a saline-paired environment. Together, these findings indicate that ACC modulation of tolerance extends beyond alteration of primary nociception, consistent with reports from chronic pain patients who have undergone cingulotomies, who state that they do not notice changes in sensation, but instead note a decrease in pain unpleasantness(41). ACC activity is also essential for conditioned placebo and nocebo responses without noxious stimuli(13, 36), as well as socially observed pain and analgesia(12). These conditioned and empathic responses reflect the anticipation of future pain and analgesia, consistent with the expectation of analgesia that occurs in the current conditioning paradigm.

Although our results demonstrate ACC’s central role in contextual analgesic tolerance, several limitations should be considered when interpreting these findings. While our approach focused on thermal nociception using the acute hot plate test, a well-established model that provides precise stimulus control and robust, reproducible behavioral responses(42), the involvement of ACC ensembles in contextual tolerance across other pain modalities remains to be determined. Future studies examining mechanical hyperalgesia, inflammatory pain, and neuropathic pain models will be essential to establish the broader clinical relevance of ACC-targeted interventions, particularly given that different pain types engage distinct ACC cell populations and circuits(19, 35). Additionally, our whole-brain imaging analysis was limited by a relatively small sample size, though the regional activation patterns observed were consistent with our independent immunohistochemistry experiments, providing convergent evidence for ACC involvement in contextual tolerance. Nevertheless, our findings provide a critical mechanistic foundation for understanding how environmental context modulates opioid analgesia.

In summary, we report a novel role for an ACC neuronal ensemble in mediating contextual opioid analgesic tolerance without affecting basal nociception. Modulating ACC activity presents a promising opportunity for rapidly reversing analgesic tolerance in environments associated with drug use, positioning targeted interventions in this region as a potential therapeutic strategy to enhance pain relief while preserving the essential ability to detect and respond to harmful stimuli.

## Methods

### Sex as a biological variable

Experiment 1 used both male and female animals. Since we did not detect statistically significant context-based tolerance reversal in female mice, subsequent experiments on tolerance reversal were performed only in male animals. Given the established sex differences in opioid effects and contextual learning, future studies will include female subjects to provide more comprehensive insights.

### Animals

All animal experiments were approved by the Yale University Institutional Animal Care and Use Committee. Male C57BL/6J mice were purchased from Jackson Laboratories. Arc- and tamoxifen-dependent Cre transgenic (ArcTRAP) mice (B6.129(Cg)-*Arc^tm1.1(cre/ERT2)Luo^*/J) were used for ensemble manipulation experiments. ArcTRAP mice were crossed in-house with tdTomato (B6.Cg-*Gt(ROSA)26Sor^tm14(CAG-tdTomato)Hze^*/J) or eYFP (B6.129X1-*Gt(ROSA)26Sor^tm1(EYFP)Cos^*/J) reporter mice to label active populations in whole-brain imaging and immunohistochemistry experiments. Mice were maintained on a 12-h light/dark cycle (lights on 7:00) with access to food and water *ad libitum*. Experiments were conducted from 10:00-18:00.

### Stereotaxic surgeries and injections

Mice (∼8-12 weeks old) were anesthetized with isoflurane (4% for induction, 1.5-1% for maintenance) in a stereotaxic frame (Kopf). Adeno-associated viral (AAVs) vectors were injected into the ACC (anteroposterior (AP) +1.00 mm; mediolateral (ML) +/− 0.35 mm; dorsoventral (DV) −1.80 mm). AAVs were injected using a manual 10 ml Hamilton microsyringe. The needle was lowered to the target, and the virus was injected at 100 nL/min. The needle remained in place for 10 min before being withdrawn. For fiber photometry studies, AAVs containing the calcium sensor GCaMP6f (pENN.AAV.CamKII.GCaMP6f.WPRE.SV40, Addgene: #100834-AAV5) were injected into the ACC unilaterally in a random order (400 nL). Optical fiber implants consisted of a zirconia ferrule (Doric Lenses) fitted with an optical fiber (200 μm core diameter, NA = 0.37), lowered to 0.1 mm above the injection DV immediately after viral injection. The ferrule was fixed to the skull using dental cement (3M). For the chemogenetic inhibition experiment, AAVs containing the fluorescent reporter mCherry (pAAV-hSyn-mCherry, Addgene: #114472-AAV5) or hM4Di (pAAV-hSyn-hM4D(Gi)-mCherry, Addgene: #50475-AAV5) were injected bilaterally (800 nL). AAV constructs were purchased from Addgene. Mice were given carprofen analgesic (5 mg/kg, intraperitoneal) and allowed to recover for at least 3 weeks before behavioral experiments. For ensemble stimulation experiments, AAV constructs containing a cre-dependent hM3Gq (pAAV-hSyn-DIO-HA-hM3D(Gq)-IRES-mCitrine, Addgene: # 50454-AAV8) or mCitrine (AAV-hSYN-DIO-HA-IRES-mCitrine, Neurophotonic, CA: # 1641-aav8) were injected into the ACC of ArcTRAP mice. Implant sites and viral expression were verified after behavioral experiments using histology. Mice that had poor or mistargeted viral expression were excluded from data analyses (n = 4). Viral expression and implant targeting in mice used in experiments is shown in **Supplementary Figure 4**.

### Conditioned opioid analgesic tolerance procedure and hotplate test of thermal nociception

Before all behavioral experiments, mice were handled and habituated to subcutaneous injections. Training occurred over 14 consecutive days with daily alternating injections. On each behavioral training day, mice were acclimated to the testing room for 1 hour before testing. Fentanyl or saline injections were randomly associated with one of two contexts with distinct multisensory features. One context consisted of a clear beaker with a flat black metal floor, was infused with an orange scent and kept in a brightly lit room. The other context consisted of a beaker lined with black and white vertical stripes and a soft gray mesh wire floor, was infused with almond scent and kept in a dimly lit room. In the initial study, both contexts were paired with saline injections and administered in an alternating, counterbalanced manner for 14 days before hotplate testing; we did not observe changes in hotplate latencies attributable to either context or repeated contextual exposure (**sFig 1)**. During training, mice received daily alternating subcutaneous injections of either fentanyl (25 μg/kg, NIDA Drug Supply Program) or saline before being placed in the respective context for 15 minutes. One day after training, antinociception and analgesic tolerance were tested by injecting mice with fentanyl and placing them in the fentanyl-associated context. Contextual tolerance reversal was tested by injecting mice with fentanyl and placing them in the saline-associated context.

Following context presentations, mice were placed on the hotplate at 56.0 +/- 0.5°C (Columbus Instruments). Mice were removed from the hotplate when nocifensive behaviors such as hind paw licking, withdrawing, flailing, and jumping were observed or after 30s (to avoid tissue damage). Mice that showed paw injury or any other type of injury before any training or testing session were excluded from data analyses (n = 6). Hot plate sessions were recorded throughout conditioning and during contextual reversal tests. Saline and fentanyl conditioning sessions were scored unblinded to ensure only mice that achieved tolerance were used in contextual tolerance tests. Contextual test sessions were scored by an investigator blinded to context, drug, genotype, or virus condition.

For the context and drug co-exposure experiment, 24 h after the last tolerance training session, mice were exposed to one of four conditions for 15 mins before hotplate testing: saline in the saline-associated context, saline in the fentanyl-associated context, fentanyl in the saline-associated context, and fentanyl in the fentanyl-associated context. 90 min after context pairings, mice were transcardially perfused, and brains were extracted and processed for immunohistochemistry.

### Whole-brain activity-dependent labeling and mapping

Male arcTRAP:Ai14 mice were injected with 4-OHT (Sigma-Aldrich) at 50 mg/kg (dissolved in a 1:9 EtOH/peanut oil solution) 1 hr after the last tolerance training session to label tolerance-active cells. 4 weeks after ensemble labeling, mice were injected with fentanyl and randomly assigned to one of two groups, one group was re-exposed to the previously fentanyl-paired context and another to the previously saline-paired context. 90 mins after the context presentation and hotplate test, mice were transcardially-perfused with ice-cold PBS containing 10 U/ml heparin, followed by 4% PFA. Brains were extracted and fixed in 4% PFA for 24 hours at 4 °C. Brains were cleared and processed by LifeCanvas Technologies using the SHIELD (stabilization under harsh conditions via intramolecular epoxide linkages to prevent degradation) protocol. Briefly, tissues were cleared for 7 days using Clear+ delipidation buffer and labeled using SmartBatch+ with rabbit anti-FOS (Abcam). Fluorescently conjugated secondary antibodies (Jackson ImmunoResearch) were applied in a 1:2 primary secondary molar ratios. Tissues were then incubated in EasyIndex (LifeCanvas Technologies) for refractive index matching (n = 1.52) and imaged using SmartSPIM. Images were registered to the Allen Brain Institute Autofluorescence Atlas and cells were quantified automatically using a neural network (LifeCanvas Technologies) as previously described(24). To determine potentially significant regions, we first devised a modified engram index previously employed by Roy *et al* (24) by dividing the mean number of activated (e.g., cFOS+) neurons in the fentanyl context-exposed group by the mean number of activated neurons in the saline context-exposed group. Regions activated in the reversal condition were determined by dividing the mean number of activated neurons in the saline context-re-exposed group by the mean number of activated neurons in the fentanyl context-re-exposed group. Brain regions with an activation index higher than 1.5 were selected for further evaluation.

### Immunohistochemistry

Mice were anesthetized with Fatal-Plus® (Patterson Veterinary) and were transcardially perfused with ice-cold PBS, followed by 4% paraformaldehyde (PFA). Brains were extracted and post-fixed overnight in 4% PFA at 4 °C. 24 hrs after, brains were transferred to a 30% sucrose solution in PBS for up to 48 hours at 4°C. Brain were embedded in Tissue-Tek® O.C.T. Compound (Sakura Finetek USA). Coronal brains sections were sliced on a sliding microtome (Leica) into at 40 μm thickness and collected into 12-well plates in PBS with 0.3% sodium azide for storage. For immunohistochemical processing, tissue was washed in PBS for 10 minutes 4 times. The tissue was then incubated with a blocking buffer consisting of 0.3% TritonX-100 in PBS (PBST) for 1 h and in PBST with 3% normal donkey serum for 1 h at room temperature. The brain sections were incubated with primary antibodies at 4 °C in a blocking buffer consisting of 1% normal donkey serum in 0.1% PBST for 24 h at room temperature. Slices were washed 4 times with PBS for 10 mins per wash and were incubated with secondary antibodies for 24 h at room temperature. After secondary antibody incubation, slices were washed with PBS 4 times and incubated with DAPI (Sigma) in PBS for 5 mins before being rinsed with PBS 4 times and being mounted onto super-frost Plus glass slides (VWR). Slices were imaged using a Fluoview FV10i microscope (Olympus) with 20x objective and 0.6 numerical aperture. Primary antibodies used for histology were: Rabbit anti-c-Fos (diluted 1:1000, Cell Signaling Technologies) and Chicken anti-GFP (diluted 1:1000, Abcam). Secondary antibodies used were Donkey anti-rabbit Alexa-Fluor 647 (diluted 1:1000, Thermo Fisher Scientific) and Donkey anti-Chicken Alexa-Fluor 555 (diluted 1:1000, Fisher Scientific).

### Fiber photometry

Mice were acclimated to the fiber photometry cable, and baseline- and foot shock-elicited calcium transients were recorded 1 week before contextual tolerance training to ensure signal quality (0.5 mA, 2 2s shocks). Mice that did not show foot shock-elicited elevations in photometric signals were excluded from further testing (n = 8). For photometry testing during contextual tolerance conditioning, mice were placed in a holding cage for 5 min. Mice were then injected with fentanyl or saline and placed in the appropriate context for an additional 15 minutes, during which signals were recorded. After exposure to the context, mice were immediately placed on the hot plate. Mice were removed from the hot plate 5 s after displaying nocifensive behaviors or after 35 s. Fluorescent signals were acquired and analyzed using custom-written MATLAB code. Signals were recorded using two LED lights at 405 and 465 nm (30 μW, Doric Lenses). The 405 channel was used as an isosbestic control, and the 465 channel was used as the calcium fluorescence channel. The first 100 s of each recording session were cropped to eliminate plugin artifacts from analysis(32). Change in fluorescence (ΔF/F) was calculated as (465 nm signal – fitted 405 nm signal)/465 signal at each time point. The Z-score for each session was calculated using the formula: Z = (x−y)/standard deviation. (where x = ΔF/F and y = mean of ΔF/F for baseline). To eliminate a plugin artifact that was present in all recordings, the first 200s of recording were clipped out of the analysis. Graphs showing the signal during context presentation were generated using the smooth data function on MATLAB with a running average of 6000. For context exposure analyses, the midpoint between the timepoint with lowest (400s) and highest (900s) ΔF/F during the first fentanyl exposure was used to separate the traces into early and late periods. Absolute signed integrals were calculated using the trapezoidal method. For hot plate signal analyses, a 7-second window centered on hot plate placement was used, with the period from 2 seconds before placement to the moment of placement serving as the pre-placement baseline, and the subsequent 5 sec as the post-placement period. To minimize transfer-induced artifacts, the signal during the transfer between the context and the hotplate (∼5s) was clipped out before analysis.

### Chemogenetic regulation of conditioned opioid analgesic tolerance

For chemogenetic inhibition studies, male C57BL/6J mice expressing either mCherry control (AAV5-hSyn-mCherry) or Gi-DREADD virus (AAV5-hSyn-hM4Di-mCherry) in the ACC underwent contextual tolerance conditioning. Mice that showed tolerance on the last day of fentanyl training were used for chemogenetic testing. 24 hrs after the last conditioning session, chemogenetic testing was initiated. On the first day of testing mice were injected with saline intraperitoneally followed 30 mins later with a subcutaneous saline injection before being placed in the saline context (saline controls). In the next session mice were intraperitoneally injected with clozapine-N-oxide (CNO; 5 mg/kg, HelloBio) followed 30 minutes later with a subcutaneous saline injection immediately before being placed in the saline context. This dose of CNO was selected based on prior studies showing a lack of sedative and locomotor effects while inducing DREADD activation (43). On the third testing session, mice received an intraperitoneal injection of saline, followed 30 mins later by fentanyl administration and placement in the fentanyl context. On the last testing session, mice were injected with CNO, and 30 mins later injected with fentanyl before being placed in the fentanyl context. For tolerance-active ensemble excitation studies, male ArcTRAP mice were stereotaxically injected with either cre-dependent mCitrine (AAV2/8-HA-DIO-mCitrine; control) or Gq-DREADD-mCitrine-containing (AAV2/8-HA-DIO-hM3Gq-mCitrine) viruses into the ACC. After 4 weeks of recovery, mice underwent contextual tolerance training. Mice that showed tolerance on the last fentanyl training session were used for further testing. 24 hr after the last training session, mice were injected with fentanyl and placed in the fentanyl context for 15 minutes before being returned to their home cage. 1 hr after context exposure, mice were lightly anesthetized and injected (Isoflurane at 1.5% at O2 200-300 cc/min for 3 minutes) with the TRAP ligand 4-OHT (50 mg/kg)(23) to prevent injection pain-induced activation as previously described(23). 3 weeks after 4-OHT injection, mice were injected intraperitoneally with CNO, 30 mins later, mice were injected with fentanyl and placed in the previously saline-paired context for 15 minutes.

### Statistics

Sample sizes for analyses are based on previously published studies that used similar approaches(13, 23, 36, 38). Data were analyzed using Prism 10 (GraphPad). All data are reported as mean ± SEM, with individual values superimposed on graphs. Statistical significance was tested using t-tests, one-way or two-way ANOVAs, and Sidak’s multiple comparisons *post hoc* test when main effects were detected. Repeated Measures ANOVAs were conducted for within-subject data when appropriate. P < 0.05 was considered statistically significant, with trends reported as exact p-values in the text.

### Study Approval

All animal experiments were conducted in accordance with the Yale University Institutional Animal Care and Use Committee (IACUC) (Protocol #2025-0789).

## Data Availability

The full dataset from this study is detailed in the main paper or the supplemental material, with raw data accessible in the Supporting Data Values file.

## Acknowledgments

We thank Nadia Jordan-Spasov and Samantha Sheppard for their technical assistance. We also thank Dr. Ralph DiLeone and Dr. Gregory J. Salimando for their conceptual assistance and manuscript discussion. Figures were partially created with BioRender. We are grateful to the NIDA Drug Supply Program for providing fentanyl for these studies.

## Author contributions

RP and MRP conceived the project and designed the studies. RP conducted behavioral, whole-brain, and photometric experiments, analyzed data, and performed immunohistochemical studies. YSM contributed to the analyses of fiber photometry and behavioral data. WZ contributed to the design of behavioral experiments. CC contributed to analyses of whole-brain and behavioral data. JT, SM, and AMT conducted immunohistochemical experiments, imaged the samples, and analyzed the data. All authors discussed the results and commented on the manuscript.

