## Supplemental Figures 1-5 for "An Anterior Cingulate Cortex Neuronal Ensemble Controls Contextual Opioid Analgesic Tolerance"

**Perez *et al.* 2026. Supplemental Figures 1-5**

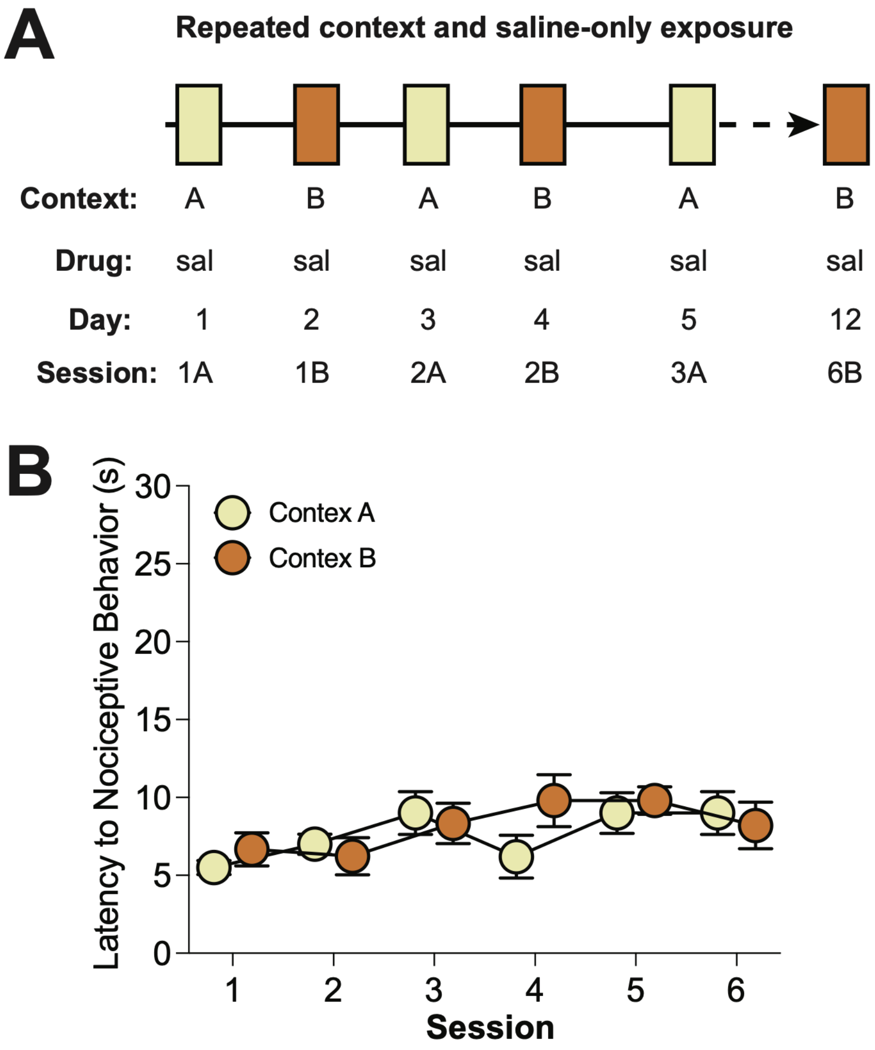


**Supplemental Figure 1. Repeated daily exposure to saline in both contexts does not alter nociceptive latencies in the hotplate test. A.** Experimental design: Male mice received alternating daily injections of saline (sal) for 12 days in distinct contexts (cx), followed by daily assessment of antinociceptive response on a hotplate at 56°C. **B.** No changes in nociception were observed after repeated sal injections (n = 10).

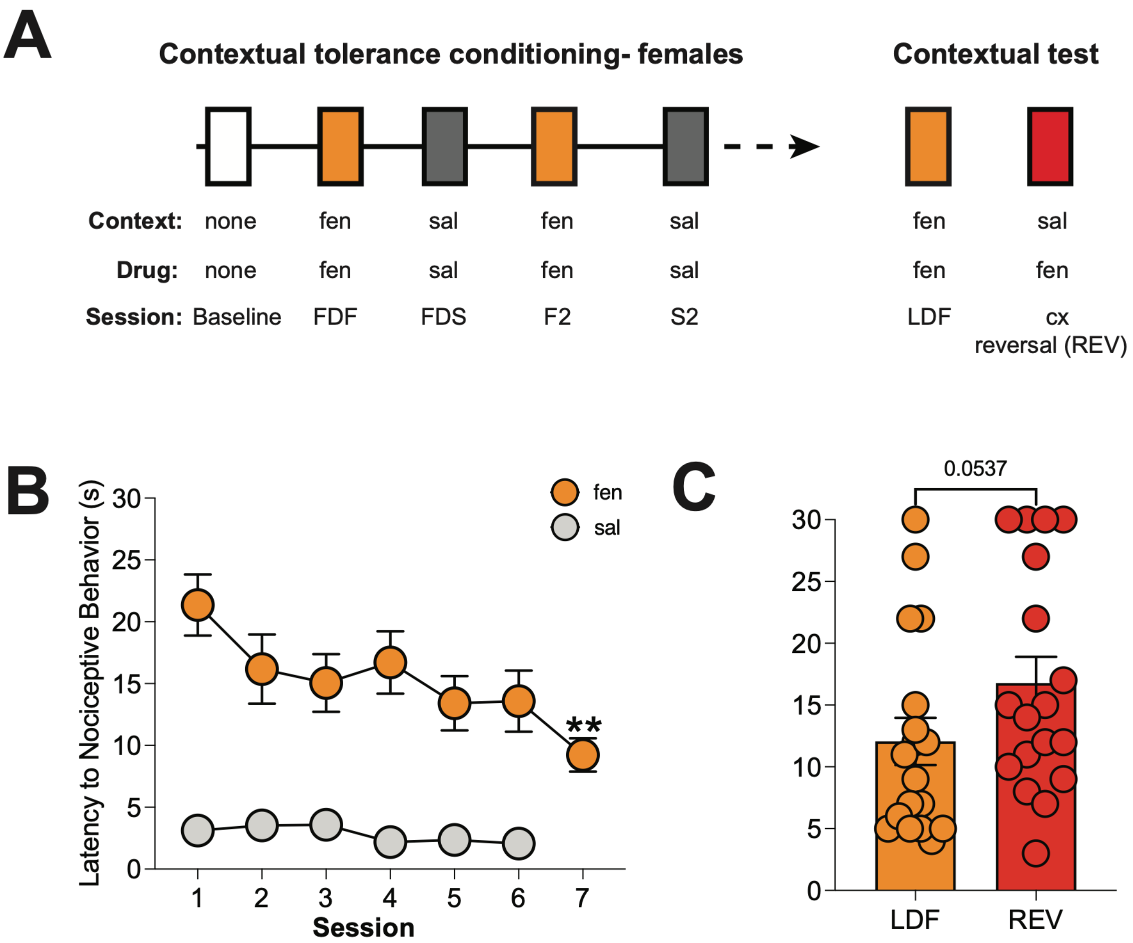


**Supplementary Figure 2. Context-dependent fentanyl administration induces analgesic tolerance in female mice and a trend towards tolerance reversal in the saline context. A.** Experimental design: Female mice received alternating daily injections of fentanyl (fen) or saline (sal) for 14 days in distinct contexts (cx), followed by daily assessment of antinociceptive response on a hotplate at 56°C. After conditioning, contextual tolerance was tested by administering fen in the previously sal-paired evaluating nociceptive latencies. **B.** A decrease in the antinociceptive effect of fen was observed over time, with a significant reduction during the 7$^{th}$ fen session when compared to the first fen session (FDF). No changes in nociception were observed after repeated sal injections. **C.** Mice showed a trend towards reversal of analgesic tolerance compared to the last day of fen administration (LDF) in the sal-paired context (REV) (n = 17).

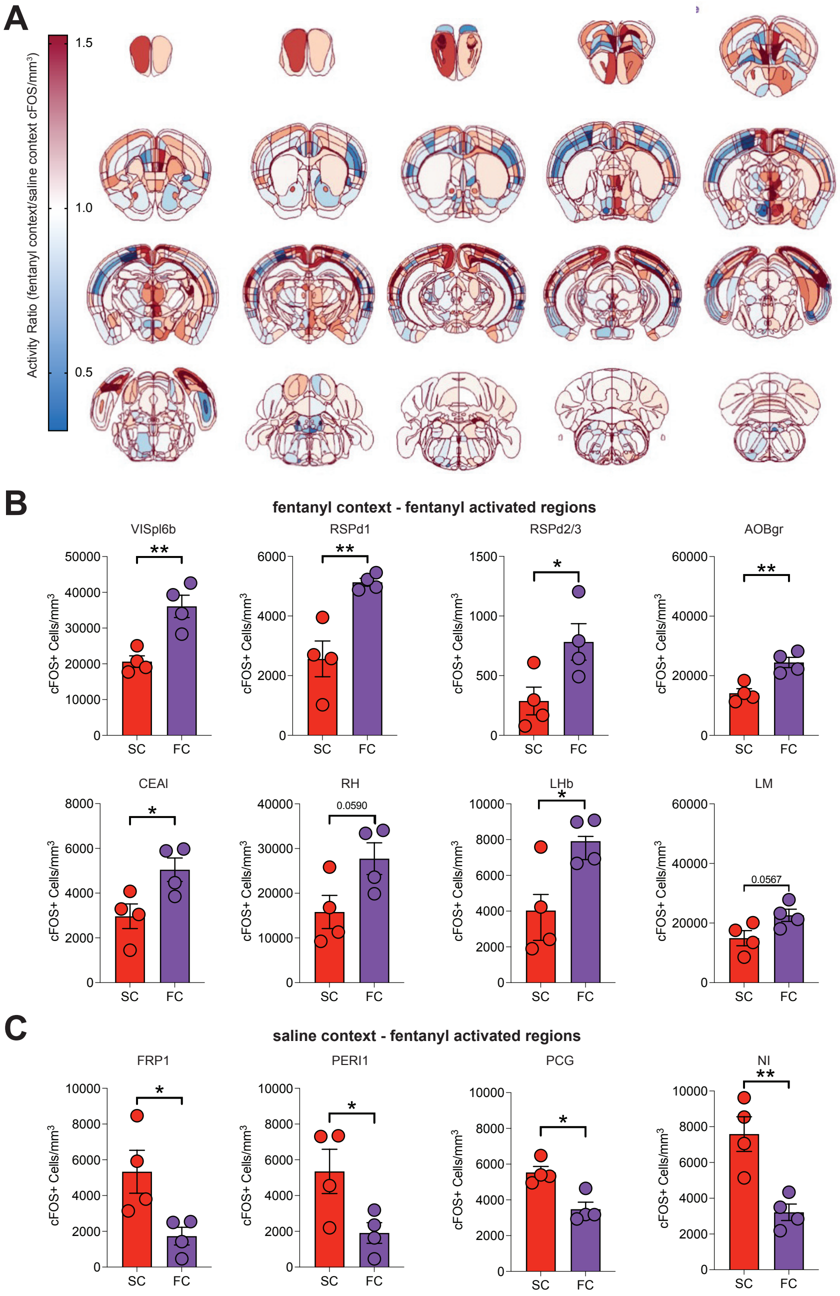


**Supplementary Figure 3. Brain regions with significantly increased cFos activity after context re-exposure. A.** Schematic showing differential cFos density across the brain, with red representing greater cFos expression following fen exposure in the fen-paired cx (FC) and blue representing greater cFos expression following fen exposure in the sal-paired cx (SC). **B.** Brain areas with increased cFos labeling following fen and fen-paired cx re-exposure. **C.** Areas with increased cFos labeling following fentanyl and sal-paired cx re-exposure (n = 4 per group; *p < 0.05, **p < 0.01). Data are shown as means ± SEM, individual data points. VISpl6b = Posterolateral visual area, layer 6b, RSPd1 = Retrosplenial area, dorsal part, layer 1, RSPd2/3 = Retrosplenial area, dorsal part, layer 2/3, AOBgr = Accessory olfactory bulb, granular layer, CEAl = Central amygdala nucleus, lateral part, RH = Rhomboid nucleus, LH = lateral habenula, LM = Lateral mammillary nucleus. FRP1 = Frontal pole, layer 1, PERI1 = perirhinal area, layer 1, PCG = Pontine central gray, NI = Nucleus incertus.

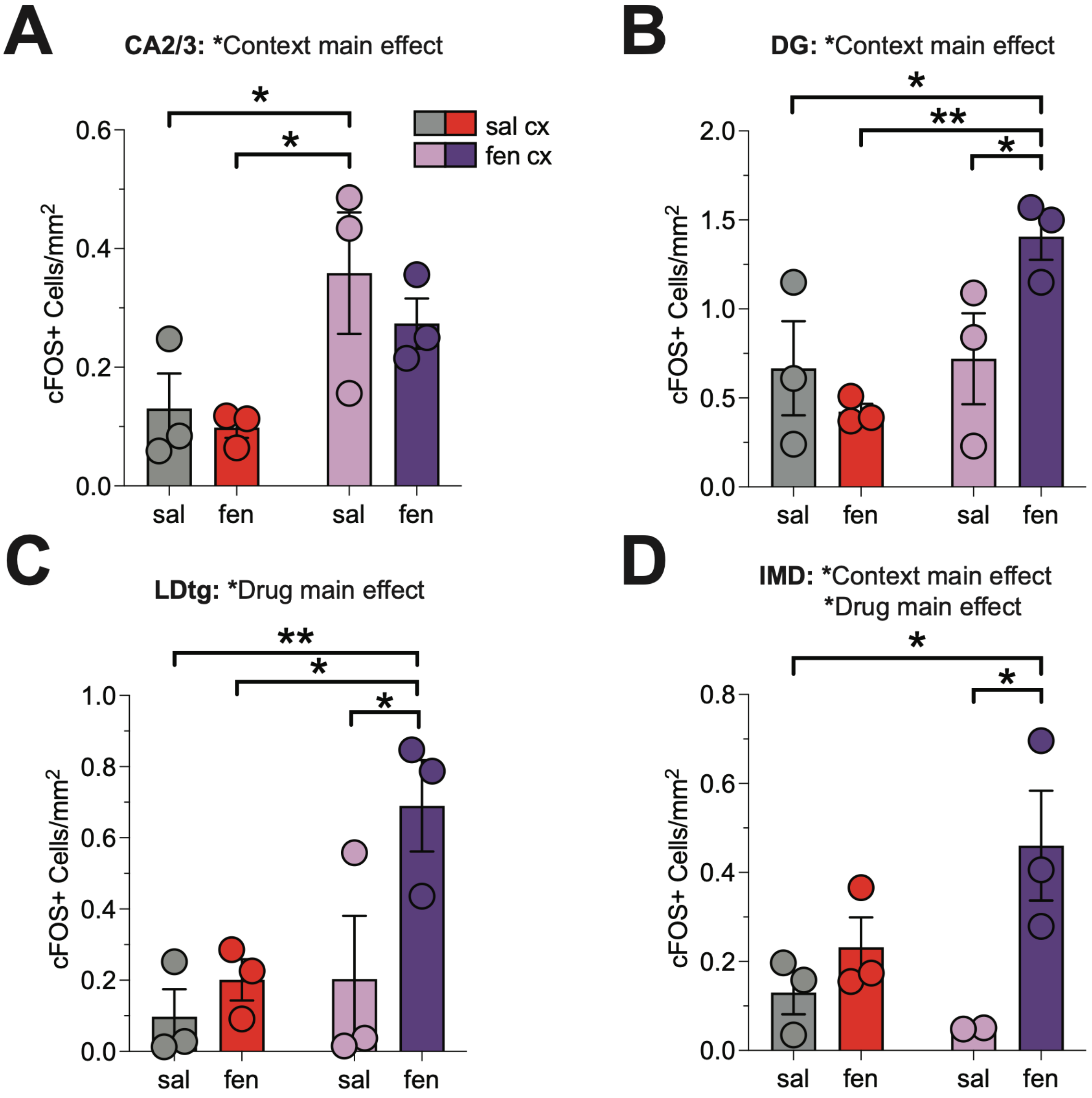


**Supplementary Figure 4. Effects of context, drug, or context and drug on cFos expression. A.** Fen-paired cx exposure in the absence of drug increased cFos labeling in the CA2/3 region of the hippocampus. **B.** cFos expression in the dentate gyrus (DG) subregion is significantly increased in response to exposure to the fen-paired cx and further increased in the cx + fen condition. **C.** There is a main effect of drug (fen exposure) on cFos labeling in the laterodorsal tegmentum (LDtg). **D.** Drug and cx have a main effect on cFos labeling in the interomediodorsal thalamus (IMD), n = 3 per group, (*p < .05, **p < .01). Data are shown as means ± SEM and individual data points.

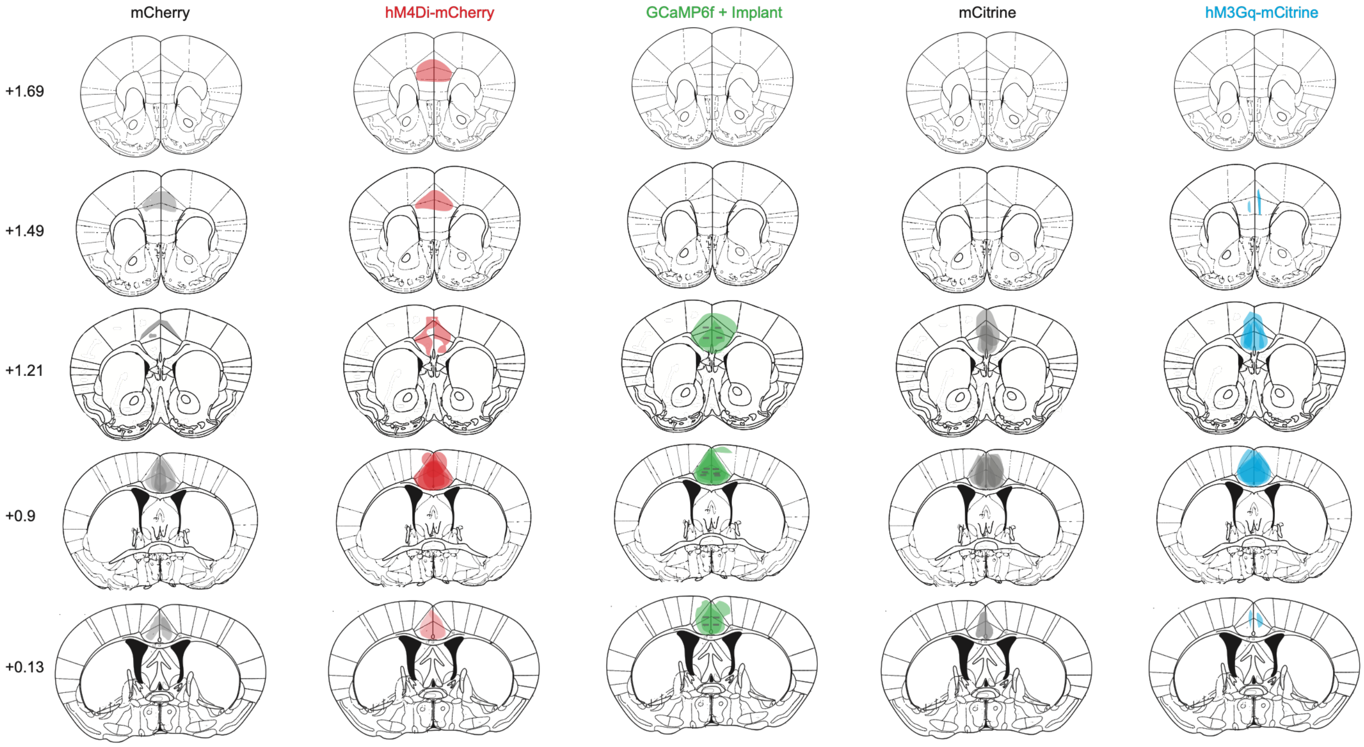


**Supplementary Figure 5. Viral expression and implant placement for all experiments. A-B.** On-target viral expression for mCherry and hM4Di in mice used in Figure 3C (2 mCherry- and 4 hM4Di mice were removed from analysis due to mistargeting or lack of viral expression). **C.** On-target GCaMP6f viral expression and bilateral implant location (hash marks) in mice used in Figure 2 (4 mice were removed from analysis due to a lack of viral expression). **D-E.** On-target viral expression for mCitrine and hM3Gq in mice used in Figure 3F (8 mCitrine- and 9 hM3Gq-infused mice were removed from analysis due to lack of viral expression).
